# FastqCleaner: an interactive Bioconductor application for quality-control, filtering and trimming of FASTQ files

**DOI:** 10.1101/393140

**Authors:** Leandro Gabriel Roser, Fernán Agüero, Daniel Oscar Sánchez

## Abstract

**Background:** Exploration and processing of FASTQ files are the first steps in state-of-the-art data analysis workflows of Next Generation Sequencing (NGS) platforms. The large amount of data generated by these technologies has put a challenge in terms of rapid analysis and visualization of sequencing information. Recent integration of the R data analysis platform with web visual frameworks has stimulated the development of user-friendly, powerful, and dynamic NGS data analysis applications.

**Results:** This paper presents *FastqCleaner,* a Bioconductor visual application for both quality-control (QC) and pre-processing of FASTQ files. The interface shows diagnostic information for the input and output data and allows to select a series of filtering and trimming operations in an interactive framework. *FastqCleaner* combines the technology of Bioconductor for NGS data analysis with the data visualization advantages of a web environment.

**Conclusions:** *FastqCleaner* is an user-friendly, offline-capable tool that enables access to advanced Bioconductor infrastructure. The novel concept of a Bioconductor interactive application that can be used without the need for programming skills, makes *FastqCleaner* a valuable resource for NGS data analysis.

## Background

The advent of Next Generation Sequencing (NGS) technologies has revolutionized genomics, transcriptomics and epigenomics research [1, 2]. The large amount of genetic information produced by these instruments requires suitable data handling and exploration methods. For most common platforms, FASTQ files are the raw starting material for subsequent analyses. A portion of the reads can include adapters or contaminants, the quality of the sequences becomes generally lower towards the end of the reads, and ambiguous base calls may be present. The correction of these and other artifacts are important steps that should be performed before using sequencing reads for mapping or assembly.

Bioconductor [3] is a widely used repository based on the R [4] programming language, containing tools devoted to the analysis of high-throughput genomic data. The massive use of these tools is, however, limited by the learning curve that users need to go through to work with customized code routines. Recently, R integration with web tools, in particular JavaScript APIs, has dramatically increased the potential of R to produce more interactive and dynamic experiences of data analysis. This integration is promissory to promote the adoption of R by many researchers for whom learning a programming language has proven to be a prohibitive investment of time and effort.

Here we present *FastqCleaner*, an R package with an offline-capable web application for QC, trimming and filtering of FASTQ files. The tool combines Bioconductor libraries for data analysis and the dynamism of a web application for data visualization.

## Implementation

### Application overview

*FastqCleaner* offers the following features:

1) Implementation of a local, offline-capable and user-friendly web interface
2) Processing of Single-Read (SR) and Paired-End (PE) files
3) Dynamic analysis of the input and output files, for customizable sampling size of reads
4) Interactive, dynamical exploration and visualization of the data, using cutting-edge technology based on JavaScript and CSS3
5) Cross-platform (running in Linux, Mac-OSX and Windows)
6) Open source, under GNU GPL (>= 2) license

### Program architecture

*FastqCleaner* was developed in R and is distributed as an R package. Data processing is controlled via R functions, that can be also accessed as normal functions from the R console (Additional file 1). These programs make extensive use of the Bioconductor packages *IRanges* [5], *Biostrings* [6] and *ShortRead* [7]. For speed improvement of the routines, C++ code was implemented in R using the *Rcpp* API [8]. The web interface included in the package was developed with *Shiny* [9], using JavaScript code written via the *jQuery* API, and CSS3.

### Design

*FastqCleaner* takes compressed or uncompressed SR or PE files as input (Fig. 1). It accepts files with qualities in both Phred+33 and Phred+64 encoding, detecting Sanger, Solexa and Illumina 1.3+, 1.5+, and > 1.8+ formats. Input files can be processed through a set of independent filters based on either one of the following two principles: 1) *Remotion of a subset of reads that do not meet a given criterion*. This group of filters can remove: a) reads with unknown bases (Ns), b) low complexity sequences, c) duplicated reads, d) reads with length below a threshold quality value, and e) reads with an average quality below a threshold value. 2) *Trimming of individual reads*. This group of filters can trim: a) full and partial adapters, b) 5′ regions below a predefined quality threshold, and c) 3′ or 5′ regions for a fixed nucleotide length. The adapter trimming algorithm extends the methodology of the *trimLRPatterns* function of *Biostrings,* designed to trim on the flanks of reads. For this purpose, *FastqCleaner* includes the *adapter_filter* function, a wrapper of *Biostrings* matching tools. The function is able to trim both adapters present on the flanks or within reads (Fig. 2). Several parameters can be passed to modify the behavior of the tool. These parameters allow, for example, to select a different threshold for the number of mismatches, to take into account the presence of indels, etc.

**Fig. 1.**
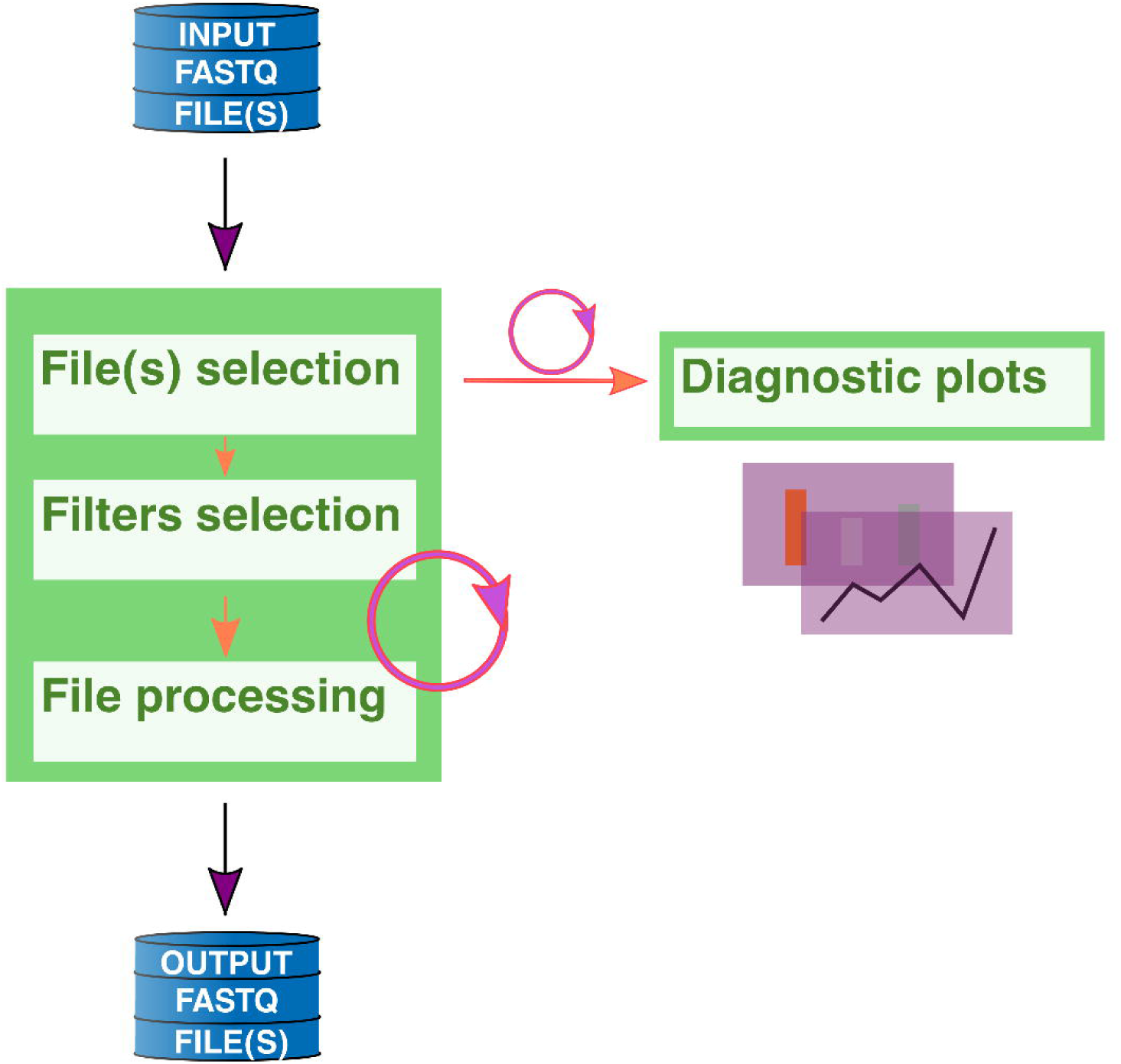
Graphical representation of a typical workflow with *FastqCleaner*, showing the initial selection of FASTQ file(s), processing, and generation of output(s). Diagnostic interactive plots can be constructed for both input and output files. Circular arrows indicate halfway points in the workflow, where different configurations can be selected to re-run the program from there.

**Fig. 2.**
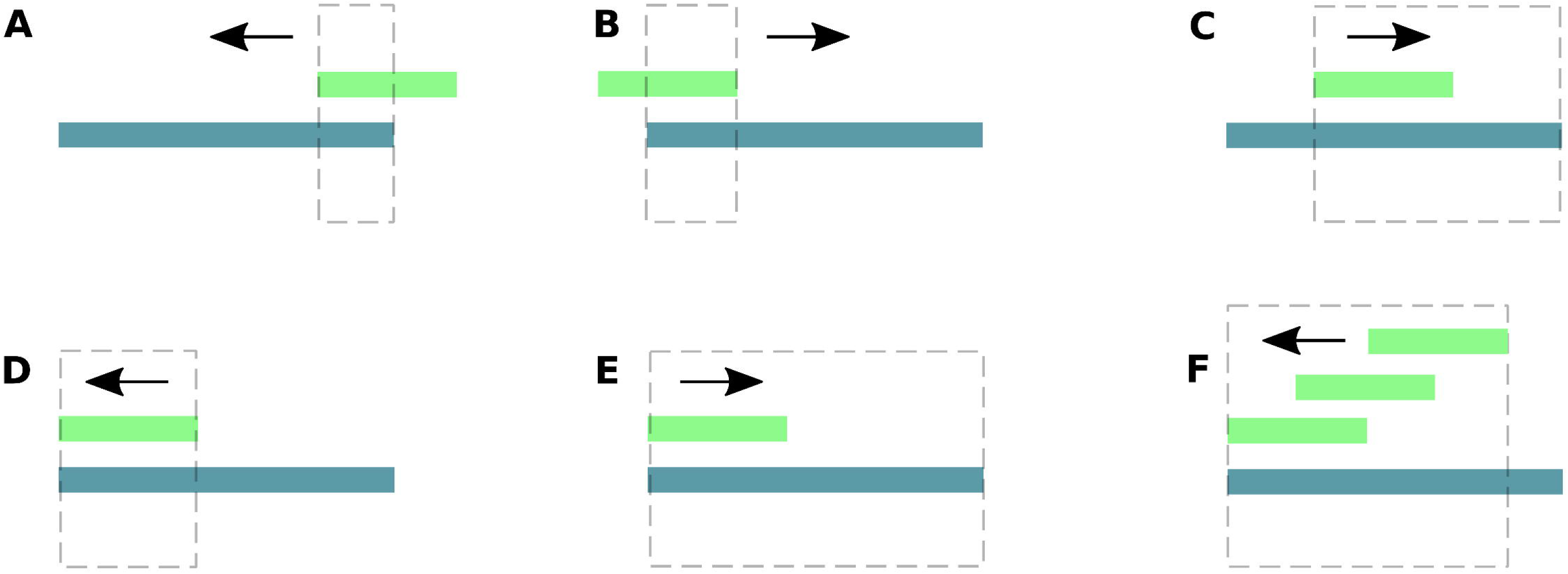
Examples for adapter trimming. Pictures show the relative position of an adapter and a read, and the expected result after processing with the *adapter_filter* function of *FastqCleaner*. Dotted lines indicate the portion of the read that will be removed. Arrows show the direction along the read used for the program to seek for matches. If one or more matches are found, the function trims the longest subsequence, that contains the matching region plus the rest of the read, in the corresponding trimming direction. *A*: partial adapter on the right + right-trimming of anchored adapter. *B*: partial adapter on the left + left-trimming of anchored adapter. *C*: partial adapter within read + right-trimming. *D,E*: full match between an adapter and a portion of the read + left-(*D*) or right-(*E*) trimming. *F*: multiple matches for a same adapter + left-trimming.

For SR files, *FastqCleaner* sequentially processes a block of reads and writes the resulting post-processed block into the corresponding output file. For PE files, the program uses in each cycle a two-step procedure: first, a block of forward and another of reverse reads are separately processed as in the SR case, and then only those reads present in both post-processed blocks are written into the corresponding output files.

### Availability

The application and a tutorial are available in Bioconductor at http://bioconductor.org/packages/FastqCleaner/

### Installation

The application can be installed following the instructions detailed at http://bioconductor.org/packages/FastqCleaner/

### Launching the application

The application can be launched with the following commands in the R console:

`library(“FastqCleaner”)`

`launch_fqc()`

Optionally, when the application is used in RStudio (versions 0.99.878 or higher), a button that allows the direct launch of the application with a single click can be found in the addins menu (Fig. 3).

**Fig. 3.**
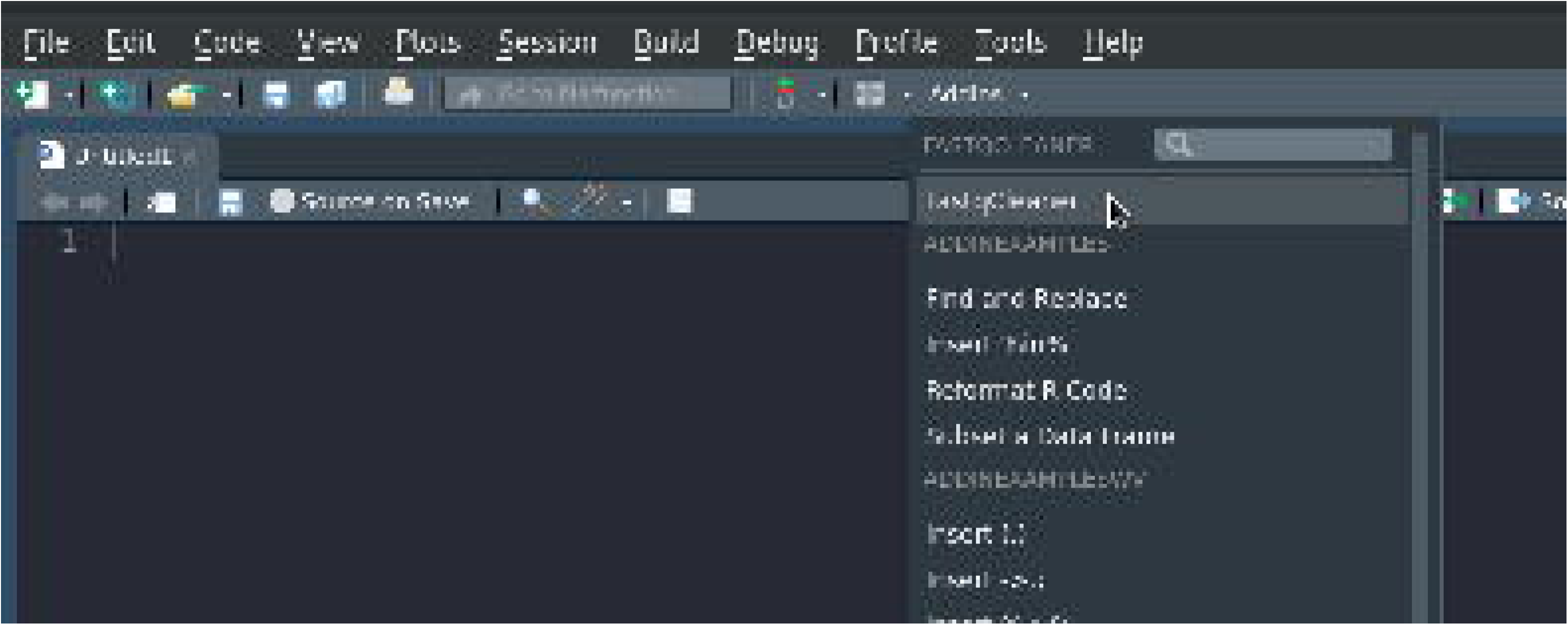
RStudio addins menu, showing the button to launch the *FastqCleaner* application.

## Results and Discussion

The web interface with its three main tabs is described in Fig. 4. The first tab (Fig. 4A) shows the file-selection menu, the available filters, and the run/reset buttons. The selection of files and filters represents the starting point in the *FastqCleaner* workflow. The second tab (Fig. 4B) shows the sequential operations performed on reads after processing. This information consists in the names of the input and output files, and a summary of informative statistics of the reads that passed the filter. The third tab (Figs. 4C, D) shows tables and interactive plots for data diagnostic. Plots can be constructed for both input (original data) and output (post-processed) files. A table with the most frequent k-mers can also be visualized.

**Fig. 4.**
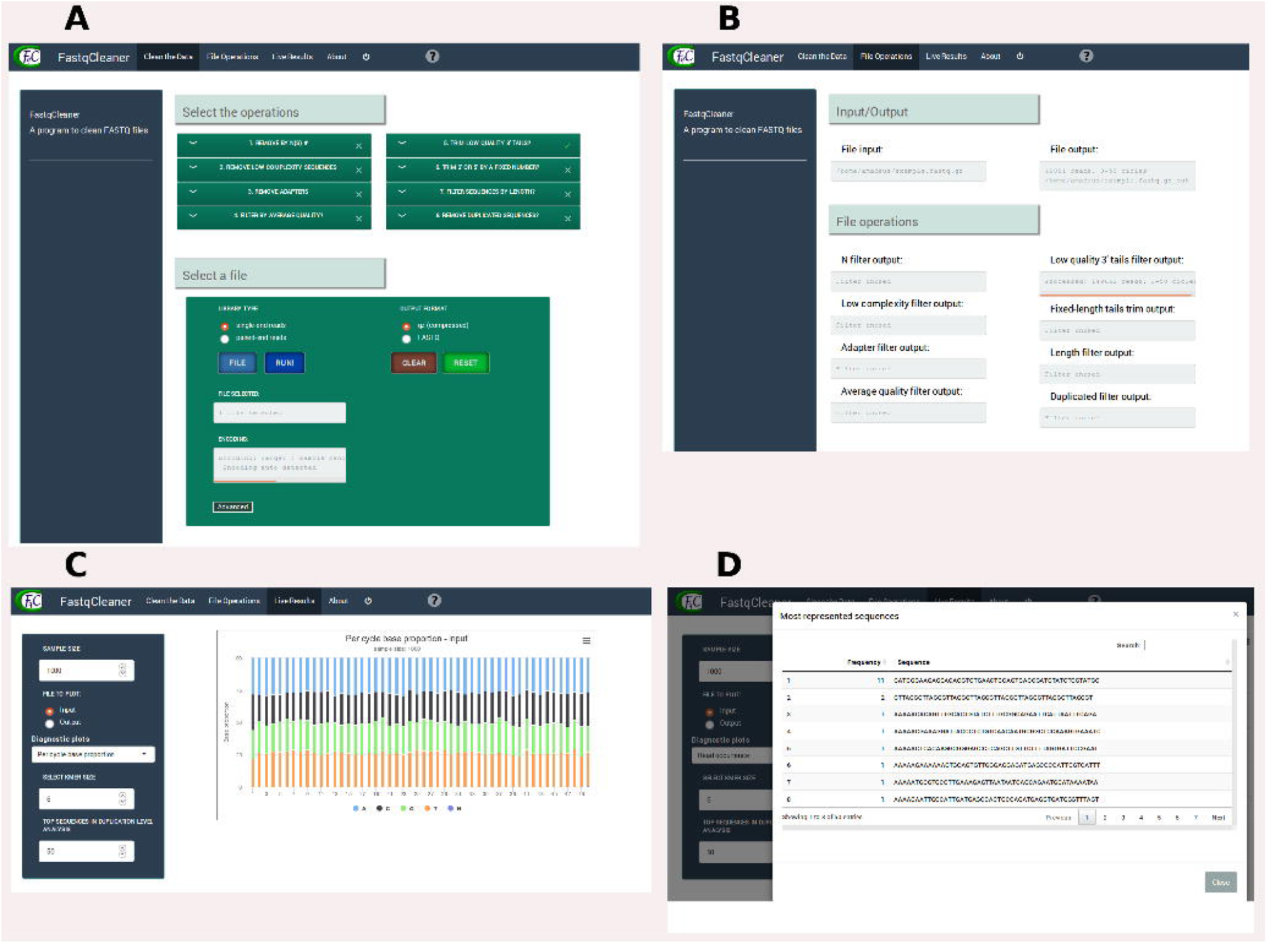
Web interface of the *FastqCleaner* application. *A*: first tab, showing an example where a file and a filter are selected. *B*: second tab, showing the processes performed after running the program. *C*: third tab, showing the analysis of the data, in this case for the input FASTQ file. The plot shows the base composition of the sequences. *D*: fourth tab, showing a table with the frequency and the sequence of each duplicated read.

A comparison of the package with other applications is shown in Table 1. Benchmarking results indicated an excellent performance of *FastqCleaner* in comparison with other pre-processing tools in terms of elapsed time. Analysis of SR pre-processing (Fig. 5) showed that these tools can be divided into three groups, in function of significant differences observed in processing speed for routine operations (Tukey HSD test, p < 0.001 for all the three pairwise comparisons). The slowest were *cutadapt* and *FASTX-Toolkit* (group 1), while *AdapterRemoval* and *Trimmomatic* (group 2) were the fastest. *FastqCleaner* showed an intermediate performance, comparable to *Skewer* and *FLEXBAR* (group 3). In PE mode, benchmarking of PE pre-processing operations (Fig. 6) showed that *FastqCleaner* significantly outperforms all other tools for routine operations (Tukey HSD test, *p* < 0.001 for pairwise comparisons of *FastqCleaner* vs other individual applications).

**Fig. 5.**
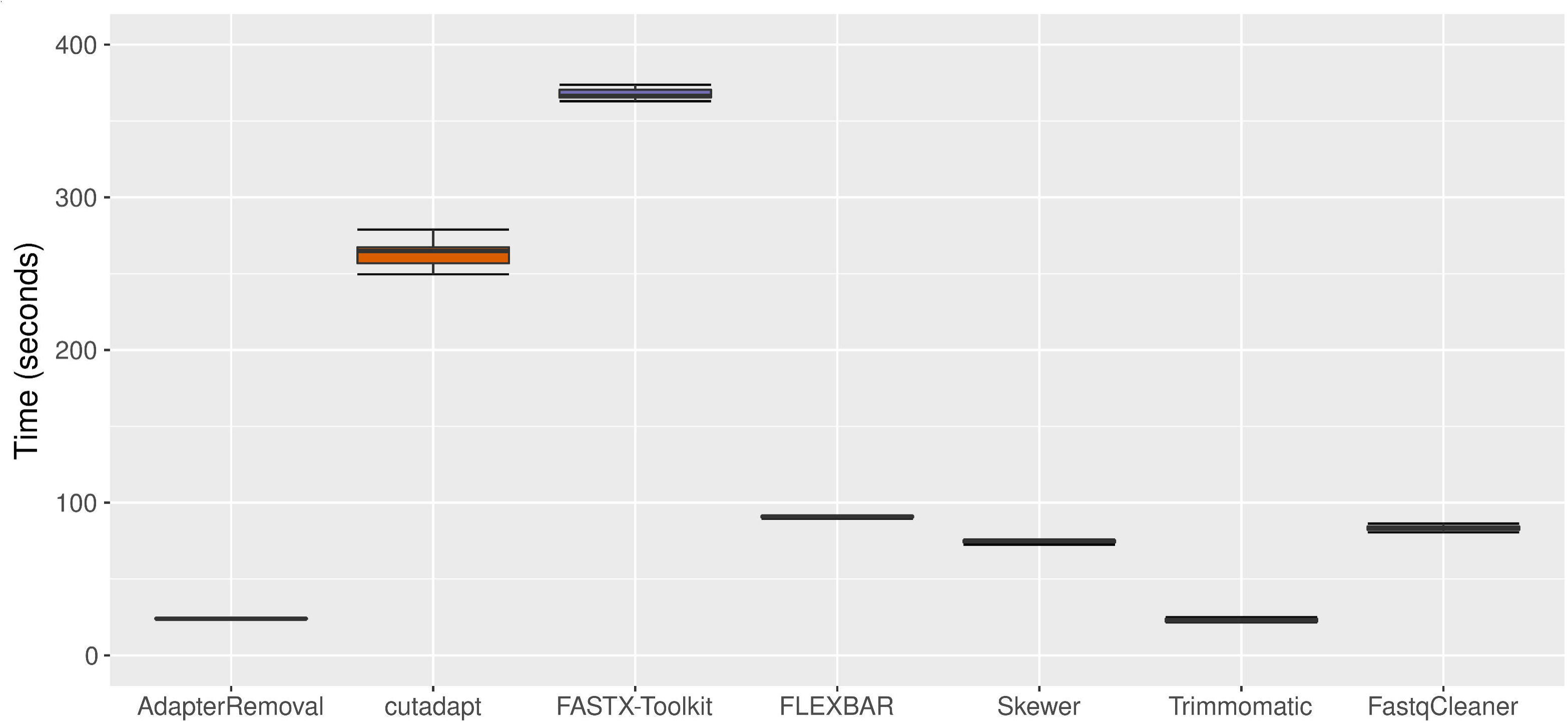
Boxplots for elapsed time (in seconds) for SR adapter trimming and read length filtering.

**Fig. 6.**
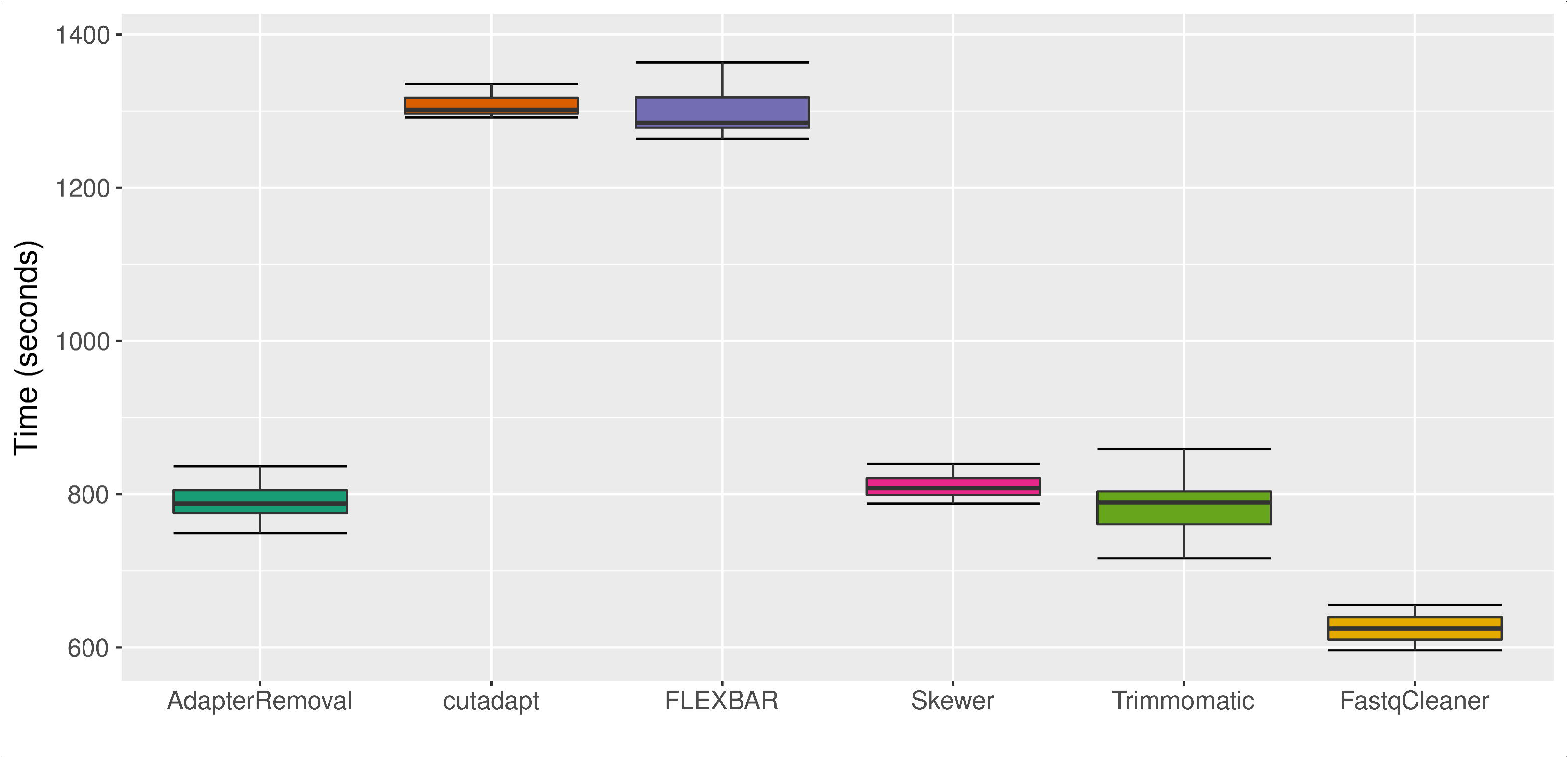
Boxplots for elapsed time (in seconds) for PE adapter trimming and read length filtering. FASTX-Toolkit is not capable to pre-process PE reads, and hence it is not shown in the plot.

## Conclusions

*FastqCleaner* is a tool with a rich and interactive cutting-edge graphical interface for pre-processing and exploration of SR and PE FASTQ files. Comparison with other available programs in a typical preprocessing scenario of adapter trimming and length filtering, showed an excellent performance of the application for both SR and PE real datasets. The application is made available as an open source license. Coding experience is not required for its use, and is therefore particularly useful for users who are unfamiliar with R programming. Furthermore, all processing happens locally in the user’s computer (even if the computer is disconnected from the network), making *FastqCleaner* amenable to run in environments where data confidentiality prevents uploading of files to the cloud.

In essence, *FastqCleaner*’s dual capability facilitates both access to the underlying state-of-art Bioconductor infrastructure and to dynamic graphical visualizations in a 100% client-side friendly web environment. This makes *FastqCleaner* a novel technological advance for the analysis of Next Generation Sequencing data.

## Methods

In order to assess the performance of *FastqCleaner*, we have compared the package with other available pre-processing tools in benchmark tests: *AdapterRemoval* 2.2.2 [10], *cutadapt* 1.14 [11], *FASTX-Toolkit* 0.0.13 [12], *FLEXBAR* 3.0.3 [13], *Skewer* 0.2.2 [14] and *Trimmomatic* 0.36 [15]. The tests (Additional file 2) were conducted for adapter removal and length filtering using SR and PE files, with 22 replicates of each tests for statistical analysis of performance. Processing conditions were standardized by disabling compression of output files and using a single thread. In addition, pre-processing in *FastqCleaner* was performed using a chunk size of 10,000 reads per cycle. For SR processing, we downloaded from SRA the dataset SRR014966, with 14.3 M reads of 36 bp. For PE processing, we downloaded the dataset SRR330569 with 27 M reads of 101 bp. Benchmark tests were conducted in R using a laptop with Linux, a 2.20GHz Intel Core i7 CPU and 16GB of 1600MHz RAM (Additional file 2).

## Additional files

Additional file 1: PDF version of the online tutorial.

Additional file 2: R script used in this work for benchmark testing.

Additional file 3: Source code of *FastqCleaner* (zip file)

## List of abbreviations

NGS: Next Generation Sequencing
SR: Single-Reads
PE: Paired-End Reads

## Availability and requirements

**Project name:** FastqCleaner

**Project home page:** http://bioconductor.org/packages/FastqCleaner/

**Operating system(s):** Platform independent

**Programming language:** R, C++, HTML, JavaScript, CSS3

**License:** GNU GPL (>= 2)

## Declarations

### Ethics approval and consent to participate

Not applicable.

### Consent for publication

Not applicable.

### Availability of data and materials

*FastqCleaner* is freely available from its Bioconductor home page at http://bioconductor.org/packages/FastqCleaner/ under a GNU GPL (>= 2) license. *FastqCleaner* can be launched on any system that has R installed. An online tutorial is available at the package home page. A PDF version of this tutorial is included as supplemental material (Additional file 1).

### Authors’ contributions

LGR designed and developed the R package. FA contributed to the improvement of the original design. LGR, FA and DOS wrote the manuscript and tested the package. DOS and FA supervised the project. All authors read and approved the final manuscript.

## Acknowledgements

FA and DOS are Career Investigators from CONICET (Consejo Nacional de Investigaciones Científicas y Técnicas, Argentina).

## References

1. Koboldt D, Steinberg K, Larson D, Wilson R, Mardis E. The next-generation sequencing revolution and its impact on genomics. Cell 2013; 155: 27–38.

2. Tripathi R, Sharma P, Chakraborty P, Varadwaj P. Next-generation sequencing revolution through big data analytics. Front Life Sci. 2016; 9: 119–149.

3. Huber W, Carey V, Gentleman R, Anders S, Carlson M, Carvalho B, Bravo H, Davis S, Gatto L, Girke T, Gottardo R. Orchestrating high-throughput genomic analysis with Bioconductor. Nat Methods. 2015; 12: 115–121.

4. R Core Team. R: A Language and Environment for Statistical Computing. R Foundation for Statistical Computing, Vienna, Austria. 2017.

5. Lawrence M, Huber W, Pages H, Aboyoun P, Carlson M, Gentleman R, Morgan M, Carey V. Software for Computing and Annotating Genomic Ranges. PLoS Comput Biol. 2013; 9:e1003118.

6. Pagès H, Aboyoun P, Gentleman R and DebRoy S. Biostrings: String objects representing biological sequences, and matching algorithms. R package version 2.44.2. 2017.

7. Morgan M, Anders S, Lawrence M, Aboyoun P, Pages H, Gentleman R. ShortRead: a Bioconductor package for input, quality assessment and exploration of high-throughput sequence data. Bioinformatics 2009; 25: 2607–2608.

8. Eddelbuettel D, Romain F, Allaire J, Chambers J, Bates D, Ushey K. Rcpp: Seamless R and C++ integration. J Stat SoftW. 2011; 40: 1–18.

9. Chang W, Cheng J, Allaire JJ, Xie Y, McPherson J. shiny: Web Application Framework for R. R package version 0.14.2. 2016. https://CRAN.R-project.org/package=shiny

10. Schubert M, Stinus L, and Ludovic O. AdapterRemoval v2: rapid adapter trimming, identification, and read merging. BMC R Notes. 2016; 9: 88.

11. Martin, M. (2011) Cutadapt removes adapter sequences from high-throughput sequencing reads. EMBnet journal. 2011; 17:10.

12. Hannon G. FASTX Toolkit. http://hannonlab.cshl.edu/fastx_toolkit/index.html. Accessed September 5 2017.

13. Dodt M, Roehr J, Ahmed R, Dieterich C. FLEXBAR—flexible barcode and adapter processing for next-generation sequencing platforms. Biology 2012; 1: 895–905.

14. Jiang H, Lei R, Ding SW, Zhu S. Skewer: a fast and accurate adapter trimmer for next-generation sequencing paired-end reads. BMC bioinformatics 2014; 15: 182.

15. Bolger A, Lohse M, Usadel B. Trimmomatic: a flexible trimmer for Illumina sequence data. Bioinformatics 2014; 30: 2114–2120.

